# Cross-Infectivity of *Vorticella* across Genera of Mosquitoes for Development of Biological Mosquito Control Strategies

**DOI:** 10.1101/822536

**Authors:** Shelby Durden, Anthony Cruz, Wayne B. Hunter, Mustapha Debboun, Dagne Duguma

**Affiliations:** University of Florida/ IFAS, Florida Medical Entomology Laboratory, Vero Beach, Florida, USA; United States Department of Agriculture, Agricultural Research Service, U.S. Hort. Res. Lab., 2001 South Rock Road, Fort Pierce, Florida, 34945. USA.; Harris County Public Health, Mosquito and Vector Control Division, Houston, Texas, USA

**Keywords:** *Vorticella*, biocontrol, *Culex nigripalpus*, *Culex quinquefasciatus*, *Aedes albopictus*

## Abstract

**P**rotozoans in general comprise about one-third of the parasitic species infecting arthropod vectors, the role of free-living ciliates on mosquitoes have been insufficiently studied either due to their low pathogenicity or being facultative parasites. Studies have shown that exposure of Paramecium ciliate protists, like *Vorticella* species, to first instar Cx. nigripalpus larvae delayed larval development and reduced biomass of emerged adults due to competition for food sources like bacteria and other microbes essential to mosquito growth and survival. Thus, we report on the capacity of a *Vorticella* protozoan’s ability to cross-infect host species and parasitize multiple mosquito larvae. The unique adapted behavior with the ability to remain on the exuviae in tree hole habitats provides a novel delivery system to develop products for target species-specific mosquitocides, larvicides, or viricides to be applied and sustained in aquatic systems.

## Introduction

Millions of deaths occur annually due to mosquito-borne illnesses such as malaria with the heaviest burden occurring in sub-Saharan Africa (WHO 2014a). Increasingly, emergence and reemergence of mosquito-borne viral illnesses such as Dengue, West Nile virus (WNV), and Zika virus epidemics, around the world are occurring (Gubler 2002; WHO 2014a,b; Liu and Zhou, 2017). To reduce the mortality and economic losses caused by mosquito-borne diseases, there is a high demand for new methods of mosquito control. Current mosquito control involves the use of chemicals and biopesticides that target the larval and adult life stages of the mosquito. These chemicals can range from organophosphates, insect growth regulators, or pyrethroids (Benelli 2015). However, past and current chemicals in use are having unintended negative effects on the ecosystem such as the emergence of chemically-resistant mosquito populations. Resistance within the population reduces the already diminished number of pesticides available for effective mosquito control (Brogdon and McAllister 1998). Stahl (2002) also compiled a report that presented the negative health and environmental risks of the four common pesticides used for mosquito control: Scourge, Anvil, Permethrin, and Malathion. Therefore, there is a renewed focus for alternative, more effective and environmentally friendly control strategies, including use of biocontrol agents or strategies which provide two avenues of attack: directly reduce mosquito populations or improve the efficacy of existing pesticides against mosquitoes.

During a field study of microbial and mosquito community dynamics, larvae of *Aedes* aegypti (L.), *Ae. albopictus* (Skuse), *Culex nigripalpus* Theobald and *Cx. quinquefasciatus* Say (Diptera: Culicidae) collected from various habitats, including artificial containers, tree holes, and waterways in Florida were found to be infected with species of *Vorticella* (Duguma et al. 2017). Over the course of four months during winter and early spring, *Vorticella* infectivity rates of mosquito larvae were found to vary across time and the type of substrate found in the larval habitat in Florida (D. Duguma, *unpublished data*). *Vorticella* is a genus of ciliates commonly found in aquatic habitats in association with mosquito larvae and other zooplankton. This ciliate protest has a contractile stalk used for attaching itself to substances *via* the means of a biopolymer glue (Cabral et al. 2017). Attachment to mobile organisms, such as the mosquito larvae, gives *Vorticella* a competitive advantage over other microbes in finding food (Kankaala and Eloranta 1987). Some studies have shown a parasitic relationship between *Vorticella* and the mosquito larval host. For example, Patil et al. (2016) demonstrated that *Anopheles stephensi* L. and *Ae. aegypti* larvae inoculated with *Vorticella* showed reduced larval growth, slower development and adult emergence. These findings indicated a potential use of *Vorticella* as a mosquito biocontrol agent to augment chemical insecticides in use. While suggested by Patil et al. (2016) that *Vorticella* may infect *Anopheles* and *Aedes* species, in this study we examined whether *Vorticella* from *Aedes* genus infects *Cx. nigripalpus* and *Cx. quinquefasciatus* and whether it impacts the two species differently. *Culex nigripalpus* and *Cx. quinquefasciatus* are prominent disease vectors in Florida and southern United States for Saint Louis encephalitis and WNV (King et al. 1960, Day and Curtis 1994, Mores et al. 2007). The larval mosquito community was examined to identify major *Vorticella* sites of attachment on the integument of the mosquitoes and whether it can transfer from larval stages to adults.

## Materials and Methods

### Mosquito collection

Twenty *Vorticella*-infected *Ae. albopictus* late (3^rd^ to 4^th^) instar larvae were collected from a tree hole located on the property of Indian River Mosquito Control District (coordinates 27.6661° N, 80.4438° W) and used as source of *Vorticella* for investigation in the laboratory. The uninfected *Cx. nigripalpus* and *Cx. quinquefasciatus* larvae were hatched from egg rafts collected from mesocosms located at the University of Florida, Institute of Food and Agriculture Sciences, Florida Medical Entomology Laboratory (coordinates 27.58672, −80.371069). Egg rafts were contained individually in wells of tissue culture plates in distilled water until they hatched. The larvae were identified to species under a dissection microscope using taxonomic keys from Darsie and Morris (2003). The larvae of each of the mosquito species were transferred to plastic trays. Larvae were fed a diet of ground brewer’s yeast and liver powder (1:1) and kept in an incubator at 27± 1 °C.

### Vorticella isolation and inoculation

The infected *Ae. albopictus* larvae collected from the field were individually dissected using sterile forceps and vortexed in one mL of 1X Phosphate Buffer Solution (PBS) solution (pH 7.2) to promote *Vorticella* suspension. The suspensions were each dispensed among two groups: Group one was: 10 *Cx. nigripalpus*/container; and Group two was: 10 *Cx. quinquefasciatus* /container. Each group had 10 plastic containers, 250 mL each, with 200 mL of deionized water. Five untreated plastic buckets each containing 10 *Cx. nigripalpus*, or 10 *Cx. quinquefasciatus* were included to serve as control treatments. The experiment was carried out in an incubator at 27± 1 °C and larval observations were made over a one month period for *Vorticella* infection. At first sighting of larval pupation, the remaining larvae were observed for infection by visual observation under a dissecting microscope, and monitored until their emergence to adults. Sixteen infected larval samples (9 *Cx. quinquefasciatus* and 7 *Cx. nigripalpus)* were placed individually in 1.0 ml of 200 proof ethanol for imaging. Light microscopy images were taken using a Keyence VHX-5000 Digital Microscope (Keyence Corporation of America, Itasca, IL, USA). Scanning Electron Microscope (SEM) images were taken using Hitachi S-4800 Scanning Transmission Electron Microscopy (STEM) (Hitachi High Technologies America, Inc., Peasanton, CA, USA) to closely examine parasitic relationship between *Vorticella* and *Ae. albopictus* larval samples. All microscopy imaging was performed at the USDA-ARS, U.S. Horticultural Research Lab, Electron Microscope Unit, Fort Pierce, FL, USA.

### Data analysis

To determine differences between *Vorticella* infection rate of *Cx. nigripalpus* and *Cx. quinquefasciatus* larvae, unpaired t-test was conducted. Non-parametric Kruskal-Wallis test followed by Dunn’s post hoc mean comparison was performed to determine differences in mortality rate between larval mosquitoes subjected to *Vorticella* and mosquitoes subjected to untreated control. A similar analysis was conducted for differences in pupation among different larval groups. The difference in total number of adults emerged between *Cx. nigripalpus* and *Cx. quinquefasciatus* larvae was evaluated using unpaired t-test. All statistical analyses and graphs were conducted using GraphPad Prism 7 (GraphPad Prism Software Inc. San Diego, CA, USA).

## Results

We examined *Vorticella’s* ability to cross-infect across genera of mosquitoes and documented the presence of *Vorticella* on different parts of larval mosquito hosts (**Fig. 1**). Using the procedure described in a previous study (Patil et al. 2016), *Vorticella* isolated from *Ae. albopictus* larvae was successfully transferred to two species of *Culex* mosquito larvae (i.e., *Cx. nigripalpus* and *Cx. quinquefasciatus*). Upon microscopic examination, major sites of infestation were determined to be along the larva’s abdominal segments, the thorax and the siphon (**Fig. 1A**). Both dorsal and ventral body parts of the second to fourth larval instars and exuviae were seen infected with *Vorticella* but not pupae or adults. *Vorticella* remained on the exuviae of the host larva during its metamorphosis to pupa suggesting that this *Vorticella* species might be restricted to immature mosquito life stages. Their attachment to the exuviae may facilitate their transition to new larval mosquito cohorts or other zooplankton.

**Figure 1.**
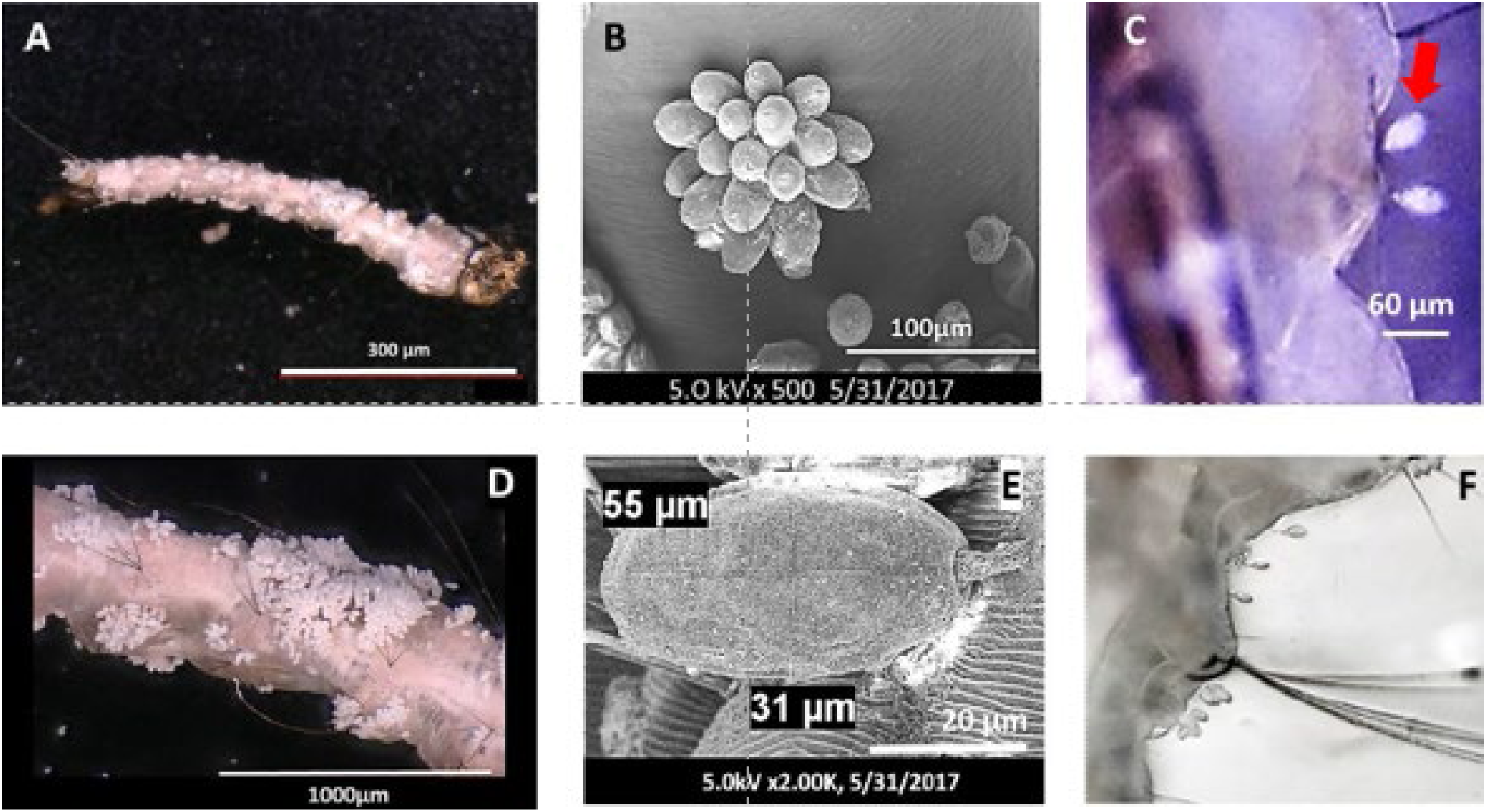
*Aedes albopictus* **(A & D)** larval samples collected from tree holes were found to be highly infested with clusters of *Vorticella*. Image **D** shows a close-up of major *Vorticella* infestation along the abdominal segments of the mosquito larva. Scanning Electorn Microsopy (SEM) images of *Vorticella* on mosquito larvae indicate polyp colony **(B)** or a singles **(C)**. A single stalk form averaged 31 µm in width and 55 µm in length **(E)**. Images of early instar *Cx. Nigripalpus* **(D)** and *Cx. quinquefasciatus* **(F)** larvae infected with *Vorticella* in the laboratory. The *red arrow* in image **C**, depicts *Vorticella* attachment as a single stalk on *Cx. nigripalpus* larvae.

The individual larvae were examined for *Vorticella* infestation at first sight of larval pupation. The infection rate differed between the two *Culex* species with significantly greater mean number of *Cx. quinquefasciatus* larvae infected with *Vorticella* than *Cx. nigripalpus* (t=3.4, df =18, *P*>*0.05*, **Fig. 2A**). On average, 52% of *Cx. quinquefasciatus* larvae were infected with *Vorticella*, whereas, only 22% of *Cx. nigripalpus* was infected with *Vorticella*. Larval mortality differed significantly between *Vorticella*-infected and non-infected mosquito larvae (Kruskal-Wallis test, *P* ≤ 0.05, **Fig. 2B**). *Culex nigripalpus* experienced greater mortality compared to *Cx. quinquefasciatus*. Greater proportion (∼20%) of *Cx. nigripalpus* larvae incurred larval mortality compared to only 10% observed in *Cx. quinquefasciatus*. Mortality in the larvae of the non-infected *Cx. nigripalpus* larvae was significantly lower than the larvae exposed to *Vorticella* suggesting that *Vorticella* may have negative impact on this species. The mean proportion of mortality rates in unexposed control (i.e., without *Vorticella* treatment) of *Cx. nigripalpus* and *Cx. quinquefasciatus* was 0 and 13%, respectively (**Fig. 2D**).

**Fig. 2.**
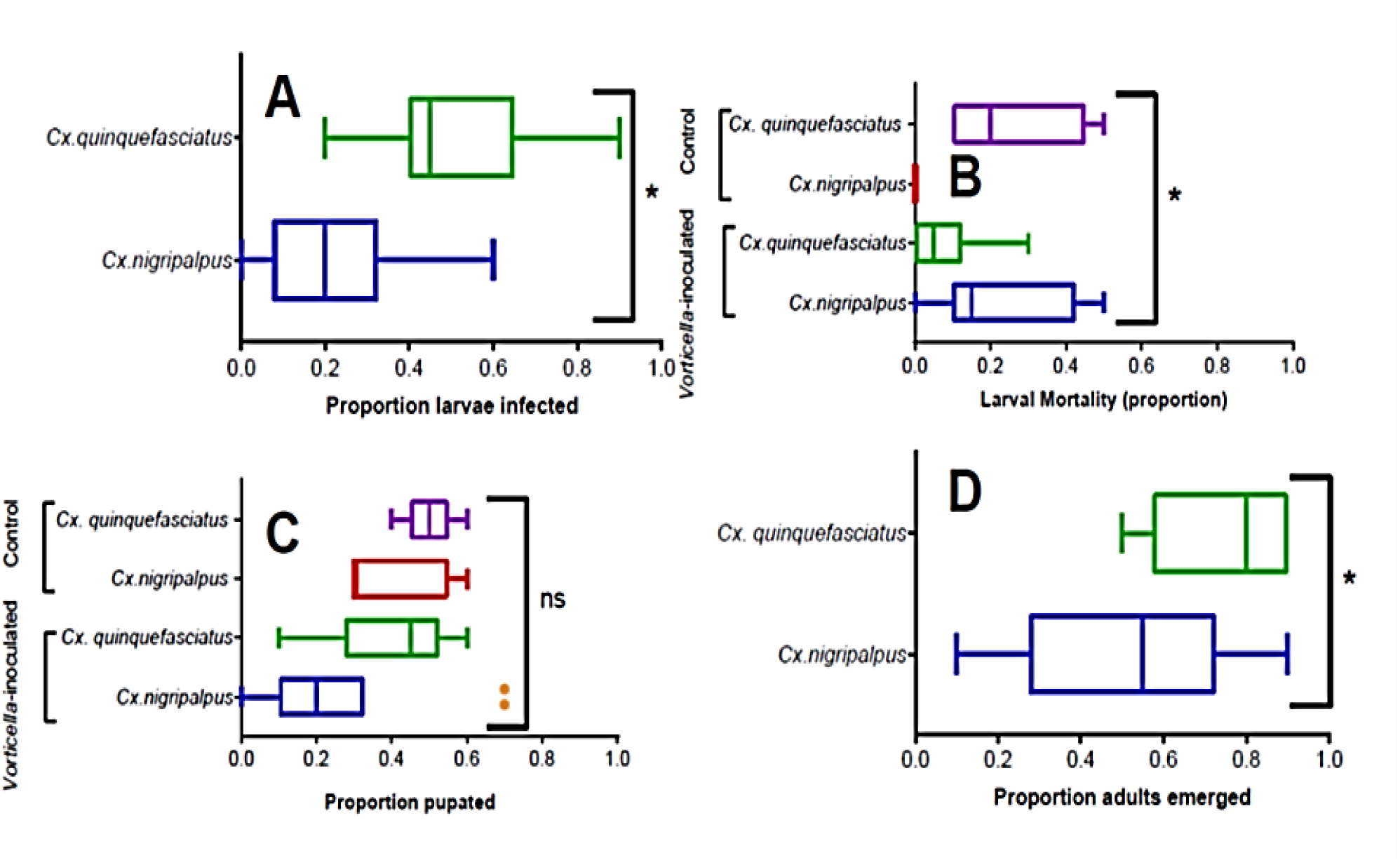
Box-and Whisker Tukey plot of mosquito *Vorticella*-infection experiment showing **(A)** infection susceptibility difference between larvae of *Cx. quinquefasciatus* and *Cx. nigripalpus*, **(B)** proportion of larvae died in both *Vorticella*-infected and control (non-infected) larvae of the two species, **(C)** proportion of mosquito larvae developed to pupae from larvae infected with *Vorticella* and control larvae, and **(D)** proportion of larvae emerged to adults following *Vorticella* infection. **\*** Indicates statistically significant differences at p ≤ 0.05, whereas **ns**= not statistically significant (p>0.05).

The difference in pupation rate among the four treatment groups was not statistically significant although slightly greater pupation rate occurred in both infected and non-infected *Cx. quinquefasciatus* and non-infected *Cx. nigripalpus* than infected *Cx. nigripalpus* larvae (Kruskal-Wallis test, *P*>*0.05*, **Fig. 2C**). However, significantly a greater number (74%) of *Cx. quinquefasciatus* emerged from the *Vorticella*-exposed larvae compared to 52% emergence rate observed in *Cx. nigripalpus* (*P* ≤ 0.05, **Fig. 2D**). These findings may suggest mosquito species-specific effects of the *Vorticella* infestation and warrant further investigation.

## Discussion

*Vorticella* is naturally occurring in freshwater environments which are also breeding sites for mosquitoes. Although the exact biological relationship is unknown, *Vorticella* appears to have a symbiotic relationship with the mosquito larvae to provide mobility to the ciliate and a competitive survival advantage. Micks’s (1955) study showed stunted growth and higher mortality rates in *Vorticella*-infected *An. atroparvus* van Thiel larvae. The effects may be due to the *Vorticella* secreting products toxic to the larvae which can cause pore formation in the larvae’s bodies or that the larvae are unable to remain on the water surface for tracheal breathing due to high levels of *Vorticella* infestation (Micks 1955).

While protozoans in general comprise about one-third of the parasitic species infecting arthropod vectors (Jenkins 1964), the role of free-living ciliates on mosquitoes have been insufficiently studied either due to their low pathogenicity, or being facultative parasites. A previous study by Duguma et al. (2017) found that exposure of *Paramecium* ciliate protists to first instar *Cx. nigripalpus* larvae delayed larval development and reduced biomass of emerged adults due to competition for food sources such as bacteria and other similar-sized microbes found essential to the growth of mosquitoes (Duguma et al., 2019). In the absence of *Paramecium*, the mosquito larvae harvested bacterial and other similar small-sized organic particles more efficiently than when found in association with the protists. Other studies have shown a severe competition for food between Vorticellid epibionts and Daphinids (Kankaala and Eloranta 1987). In addition, the heavy infestation of *Vorticella* hampers mobility of its host subjecting the host susceptible to predation (Kankaala and Eloranta 1987). Although we have not measured growth performances of larvae in our study, the higher mortality observed in *Cx. nigripalpus* larvae may be attributed to competition by *Vorticella* for bacterial food sources in addition to their possible physiological effects on the larvae. However, a similar effect was not observed in *Cx. quinquefasciatus* suggesting that their effect might be species-specific. These findings show promise in the utilization of ciliates in mosquito population control.

The protozoan’s ability to cross-infect and parasitize multiple mosquito larvae and its ability to remain on the exuviae provides a unique delivery system for novel species-specific mosquitocides, or viricides to be applied and sustained in aquatic systems. The need for studies to evaluate these advantages of *Vorticella* as a potential delivery system, as a form of biological control to reduce mosquitoes and the spread of vector-borne pathogens, such as Dengue, Zika and West Nile Virus are being pursued as a sustainable mechanism for mosquito control.

## Acknowledgements

We thank Maria T. Gonzalez, Senior Technician, USDA, STEM Microscopy unit, Fort Pierce, FL., Jacqueline L. Metz, technical assistance, USDA, and Matthew Hentz for instruction on the VHX-5000 digital microscope, and Dr. Joseph Krystel for his valuable comments. This study was supported by funding from Florida Department of Agriculture and Consumer Services.

*This research was supported in part by the U.S. Department of Agriculture, Agricultural Research Service.” Mention of trade names or commercial products in this publication is solely for the purpose of providing specific information and does not imply recommendation or endorsement by the U.S. Department of Agriculture. USDA is an equal opportunity provider and employer.*

